# A semi-automated machine learning-aided approach to quantitative analysis of centrosomes and microtubule organization

**DOI:** 10.1101/2020.01.03.894071

**Authors:** Divya Ganapathi Sankaran, Bharath Hariharan, Chad G. Pearson

## Abstract

Microtubules (MTs) perform important cellular functions including migration, intracellular trafficking, and chromosome segregation. The centrosome, comprised of two centrioles surrounded by the pericentriolar material (PCM), is the cell’s central MT organizing center. The PCM proteins, including *γ*-tubulin and Pericentrin, promote MT nucleation and organization. Centrosomes in cancer cells are commonly numerically amplified. However, the question of how amplification of centrosomes alters the MT organization capacity is not well-studied. We developed a quantitative image-processing and machine learning-aided approach for the automated analysis of MT organization. We designed a convolutional neural network-based approach for detecting centrosomes and an automated pipeline for analyzing MT organization around centrosomes, encapsulated in a semi-automatic graphical tool. Using this tool, we analyzed the spatial distribution of PCM proteins, the growing ends of MTs and the total MT density in breast cancer cells. We find that breast cancer cells with supernumerary centrosomes not only have increased PCM protein but also exhibit expansion in PCM size. Moreover, centrosome amplified cells have a greater MT density and more growing MT ends near centrosomes than unamplified cells. The semi-automated approach developed here enables facile, unbiased and quantitative measurements of centrosome aberrations. We show that these aberrations increase MT nucleation and promote changes to MT density and the spatial distribution of MTs around amplified centrosomes.

## INTRODUCTION

Microtubules (MTs) are cytoskeletal polymers that perform biological functions essential for life. The interphase MT array is required for cell migration, intracellular trafficking, and cell polarization while the mitotic MT array organizes the bipolar spindle and promotes faithful chromosome segregation (Desai and Mitchison, 1997; Hyman and Karsenti, 1996; Inoue’ and Salmon, 1995). Consisting of polar tubulin subunits, MT polymers have a dynamic plus end and a less dynamic minus end (Allen and Borisy, 1974; Bergen and Borisy, 1980; Desai and Mitchison, 1997; Walker et al., 1988). In cycling cells, minus ends are generally focused at the cell’s MT organizing center or centrosome where MTs are nucleated and anchored (Brinkley, 1985). Centrosomes consist of a pair of centrioles surrounded by the pericentriolar material (PCM). Pericentrin and CDK5RAP2 scaffold the recruitment of the *γ*-tubulin ring complex (γ-TuRC) which nucleates MTs (Dictenberg *et al*., 2002; Fong *et al*., 2007; Farache *et al*., 2018, Stearns and Kirschner, 1994; Moritz *et al*., 1995; Zheng *et al*., 1995). During the G1 phase of the cell cycle, cells have one centrosome and two centrioles. Centriole duplication occurs in S-phase resulting in two centrosomes, each with two centrioles (Hinchcliffe and Sluder, 2001; Piel et al., 2000; Vorobjev and Chentsov, 1982). Concurrent to the centriole duplication cycle, PCM organization occurs as a toroid around mature centrioles during interphase, and expands into a more amorphous structure in preparation for mitosis (Fu and Glover, 2012; Lawo et al., 2012; Mennella et al., 2012; Mennella et al., 2014; Sonnen et al., 2012; Woodruff et al., 2014; Woodruff et al., 2015). Centrosome duplication doubles the MT nucleation capacity during interphase (Salaycik, 2005). Therefore, the number of centrosomes and expansion of the PCM promotes the MT architecture of the cell.

Defects in centrosome number control commonly occurs in cancer cells resulting in centrosome amplification (CA) (D’Assoro et al., 2002; Denu et al., 2016; Ganapathi Sankaran et al., 2019; Guo et al., 2007; Lopes et al., 2018; Marteil et al., 2018; Salisbury et al., 2004; Schneeweiss et al., 2003). We and others previously quantified the changes in centriole number in cancer cells but, to date, how the PCM changes with more centrioles remains largely unknown. Moreover, how CA affects the cell’s MT organization during interphase is not known. This is important as breast cancer therapeutics that target MTs, including Taxol, have been used for decades (Pazdur et al., 1993; Schiff and Horwitz, 2006). This highlights the need for quantitative studies of MT organization in normal and breast cancer cells.

Tracking individual MTs within cells is complicated by high MT densities. However, MT organization and dynamics can be measured by quantifying the distribution of fluorescently labeled MTs and/or MT-plus end binding proteins, such as the End-Binding Protein 3 (EB3), which localizes to growing MT-ends (Roth et al., 2019; Semenova and Rodionov, 2007; Stepanova et al., 2003; Straube and Merdes, 2007). Counting EB3 foci, or “comets”, is time-consuming and requires quantitative computational approaches (Applegate et al., 2011). Moreover, these tools require extensive manual annotation and the detection of centrosomes. Automation of this analysis requires the automated detection of centrosomes, the identification of EB3 comets around detected centrosomes and the analysis of EB3 comet distributions. Similar approaches are also required to quantify the distribution of MTs surrounding centrosomes.

In this study, we establish a semi-automatic analysis pipeline that automatically detects centrosomes using machine learning while allowing for user-defined correction of errors. Machine learning is a class of techniques where a computational model is “trained” to emulate the annotations of an expert. The problem of detecting centrosomes introduces additional challenges to extant machine learning pipelines: the number of training annotations is too small to train existing machine learning models, and the large size of the image files makes existing models run slowly. Here, we introduce a new pipeline that tackles these challenges so that training is efficient and the run time is quick. The identification of centrosomes by machine learning is then coupled to automated analysis pipelines for the PCM, MTs and EB3 comets around the identified centrosomes. Finally, the entire pipeline is encapsulated into a new graphical tool that allows users to visualize and correct errors that might be performed by the automated analysis.

This semi-automated image analysis pipeline was used to investigate centriole and centrosome frequency, PCM size and MT distributions in normal and breast cancer cells with amplified centrosomes. We discovered that breast cancer cells with amplified centrosomes exhibit increased PCM protein levels and PCM size. Moreover, cells with amplified centrosomes have elevated MT density and MT growth near centrosomes. In summary, centrosome amplification increases MT nucleation and promotes changes to MT density and spatial distribution of MTs around amplified centrosomes in breast cancer cells.

## RESULTS

### Machine learning for centrosome detection and cell segmentation

Our first step in creating an automated analysis pipeline for centrosomes and MTs was to automate the task of centrosome detection using machine learning.

Centrosome foci can be difficult to detect because of large variations in the signal-to-noise ratio. In preliminary experiments, we found that simple image processing techniques that rely on consistent signal-to-noise ratios often failed to detect these foci accurately. An alternative algorithm for detecting centrosomes was designed by adapting existing object detection protocols using computer vision (Liu et al., 2016). This prior work uses convolutional networks^1^ to assign a score to every location in the image. The score output at a particular location is interpreted as the probability of a centrosome at that location. The convolutional network is “trained” on a dataset consisting of images with annotated locations of centrosomes.

We found that using prior out-of-the-box convolutional network architectures used in computer vision presented two major challenges. First, these models require millions of training images that are not pertinent to biological samples (He and Sun, 2014). This is because the convolutional network has a large number of parameters; with more parameters, more data are required to optimize these parameters. Second, these architectures are designed to run on small images (typical sizes are 224 x 224 pixels) and so are very slow when run on large fluorescence microscopy images (typically greater than 1000 x 1000 pixels).

To address these challenges, we designed a new convolutional network architecture that requires less training data, memory and time to run (Figure 1A). The input to the network is a 3-dimensional fluorescence image stack where each XY coordinate is projected to their maximum pixel intensity. The mean and standard deviation of the fluorescence intensity across all pixels is calculated, and the mean is subtracted from each pixel and divided by the standard deviation. Subtracting the mean makes the result invariant to changes in brightness in the original image (or more precisely, the addition of a constant to the intensity of all pixels). Analogously, dividing by the standard deviation makes the result invariant to changes in contrast (or more precisely multiplication of the intensity of all pixels by a constant). The result is thus brightness- and contrast-normalized.

**Figure 1:**
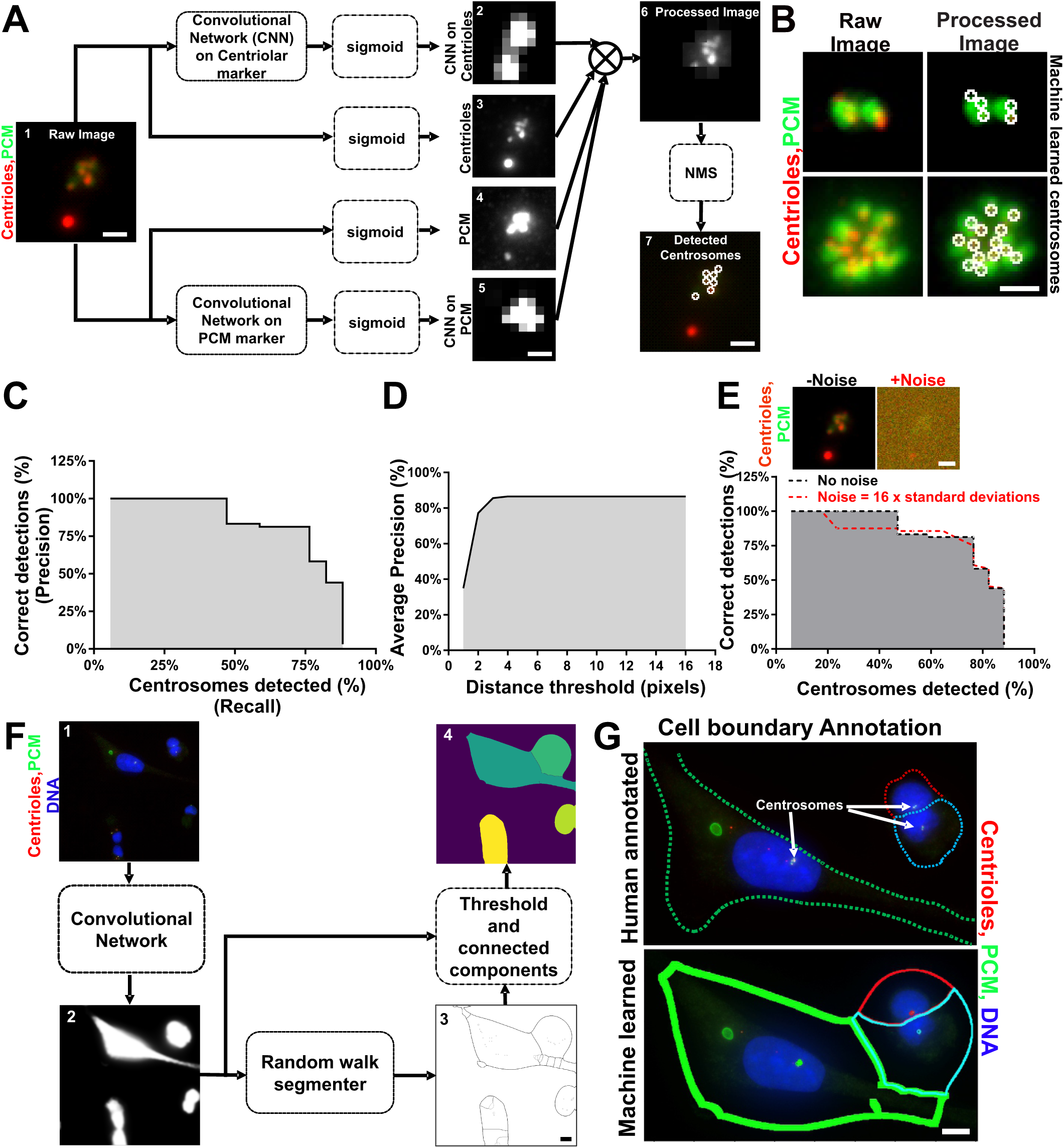
A machine learning algorithm for centrosome and cell detection. **(A)** Separate convolutional networks operating on centrin (centriole, red) and Pericentrin (PCM, green) fluorescence signals assign a score for each pixel indicating the likelihood that it is a centrosome (panels 2 and 5). These are passed through a sigmoid function (f(x) = 1/(1+e^-x^)) and multiplied together with the intensities of the centrin and Pericentrin channels (panels 3 and 4), also after a sigmoid function, to produce a final score for each pixel (panel 5). Peaks in the final score are detected using non-max suppression and identified as centrosomes (panel 7). Scale bar represents 1 μm **(B)** Left panels, raw image of centrosomes stained for centrioles (centrin, red) and γ-tubulin (green). Right panels, detected centrosomes (white). Scale bar represents 1 μm. **(C)** Evaluation of the centrosome detection in terms of the fraction of centrosome detections that are deemed correct (precision) and the fraction of centrosomes detected (recall). The plot shows precision and recall as the selection criterion is made less conservative by reducing the threshold score at which a centrosome is detected. **(D)** Variation of average precision (averaged over multiple recall values) as the criterion for a correct detection is made more lenient. The x-axis is the maximum distance between a true and a predicted centrosome at which the predicted centrosome is still considered correct. **(E)** Precision and recall values for a noisy image compared to the original image. Scale bar represents 1 μm. **(F)** Pipeline for segmenting out individual cells. 1 original image, 2 output of the neural network that identifies pixels that fall within cells, 3 boundary map obtained from the random walk segmenter with estimated boundary strength, 4 final segmentation. Scale bar represents 10 μm. **(G)** Comparison between predicted cell segmentation and human-annotated cells. Scale bar represents 10 μm.

The network then uses this normalized intensity image to compute four images (corresponding to four scores per pixel; Figure 1A, panels 2-5). Two of these images are the fluorescence intensity for centriole (Figure 1A, panel 3) and PCM markers (Figure 1A, panel 4); we use these because co-localization of high-intensity foci in these channels likely indicates centrosomes. By themselves, these channels might be noisy and produce spurious foci. We obtained two more images by running a small, fast 4-layer fully convolutional network on each individual channel (Figure 1A, panels 2 and 5). The network reduces the resolution by a factor of 8, but also reduces noise and removes spurious foci. Reducing resolution has two benefits. First, it makes the network much faster. Second, because the convolution operations used by the network operate on fixed-size image patches, at a lower resolution these patches correspond to a larger fraction of the image, thus allowing the network to look at a larger part of the image to make decisions for each pixel. All outputs are up-sampled using nearest-neighbor interpolation to the size of the original image and converted to a score between 0 and 1 using the sigmoid^6^ function (f(x) = 1/(1+e^-x^)) before being multiplied together, resulting in a final accurate score for each pixel corresponding to the likelihood of finding a centrosome at that location (Figure 1A, panel 6). The full model has fewer than 10,000 parameters and is very efficient (fewer than 7 gigaflops, compared to 35 gigaflops for a standard object detection approach; Liu et al., 2016).

Centrosomes were identified as local maxima in this output (circles in Figure 1A, panel 7). More precisely, pixels in the output were considered sequentially in decreasing order of scores. The highest scoring pixel was declared as a centrosome, and subsequent, lower scoring pixels were declared centrosomes only if no previously declared centrosome was fewer than r=5 pixels (0.65 μm) away (this is called non-max suppression;(Viola and Jones, 2011)). This produces a ranked list of putative centrosome detections, and the user can choose where to threshold this list. The full centrosome detection pipeline is shown in Figure 1A and example detections are shown in Figure 1B.

To train the model, we annotated centrosomes on a small dataset of 10 images and used these as training images to train the convolutional networks. To evaluate this centrosome detection approach and make sure that it was indeed detecting centrosomes correctly, we annotated another image that the network did not see during training. We then marked detected foci as correct if they were within 5 pixels (0.65 μm) of a centrosome, and as spurious otherwise. If multiple foci were detected where there was only one centrosome, all but one of the foci were considered spurious. We then measured the precision (the fraction of detected foci that were deemed correct) and the recall (the fraction of centrosomes that were detected). An ideal algorithm would detect all and only correct centrosomes, achieving precision and recall of 100%. We plotted how precision varies with recall as the score threshold for declaring a centrosome is reduced (Figure 1C). While not perfect, the centrosome detector maintained a precision of 75% even at high recall (>75 %). Note that the recall is not 100% because of some detections erroneously suppressed by the non-max suppression procedure.

We next evaluated the ability of the centrosome detector to accurately localize the centrosome (Figure 1D). As above, detected foci were marked correct if they were within a threshold distance of a centrosome. We varied this distance threshold and at each distance threshold, we computed the average precision, or the precision averaged over multiple values of recall. We find that the centrosome identification maintains a high average precision even for stringent thresholds (a distance of 5 pixels corresponds to 0.65 μm), indicating that the detector accurately localizes centrosomes. Finally, we asked how resilient the centrosome detection is to noise. We artificially added Gaussian noise to the image and computed the precision versus recall curve for the centrosomes identified on a noisy image (Figure 1E). Even with noise more than 16 times the standard deviation of the original image, centrosomes were consistently detected. However, note that the assumption of Gaussian noise may not correspond to the noise observed in real microscopy images.

Next, the analysis requires that individual cells are separately analyzed. To group the detected centrosomes into each individual cell, we created a pipeline for segmenting the cells in the image (Figure 1F). In the first step, a convolutional network identifies all pixels that fall inside a cell. This convolutional network is trained using a small dataset where the cells have been annotated by hand. The output of this network can be interpreted as a probability *p(x)* for each pixel *x* that indicates whether it falls inside a cell. The next step is to group the pixels with a high *p(x)* into separate cells. To do this, local peaks were identified in this output as markers for possible cells and a random walk segmenter (Grady, 2006) was used to segment the cell pixels by assigning each cell pixel to one of the markers. However, the random walk segmenter can over-segment and subdivide a single cell into multiple cells. We, therefore, estimate the strength of the boundary between the cells using the convolutional network output. We define this boundary strength as 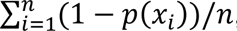, where the summation is over all pixels on the cell boundary (Figure S1A). A user-defined threshold was used to merge cells that are separated by weak boundaries. We qualitatively compared the result of the semi-automatic segmentation approach to a human-annotated cell. Although the machine-generated segmentation does not accurately track cell boundaries, it is able to identify all the cells and capture the bulk of the interior of the cell especially in the vicinity of the centrosome (Figure 1G).

Together, these results suggest that our centrosome detection and cell segmentation approach can increase the speed and accuracy of analyses but may require manual intervention when predictions are incorrect.

### Evaluation of PCM and MT organization using machine learning aided image processing

Once centrosomes and cell boundaries have been detected, we designed automated image processing algorithms to analyze the spatial distribution of centrosomal PCM proteins, MTs and EB3 comets at and around the centroid of centrosomes (Figure 2A-D). This analysis routine can be automatically run on cells with detected centrosomes. These image processing routines and the machine learning-aided detection of centrosomes and cells described above were encapsulated into a semi-automatic graphical tool for analysis.

**Figure 2:**
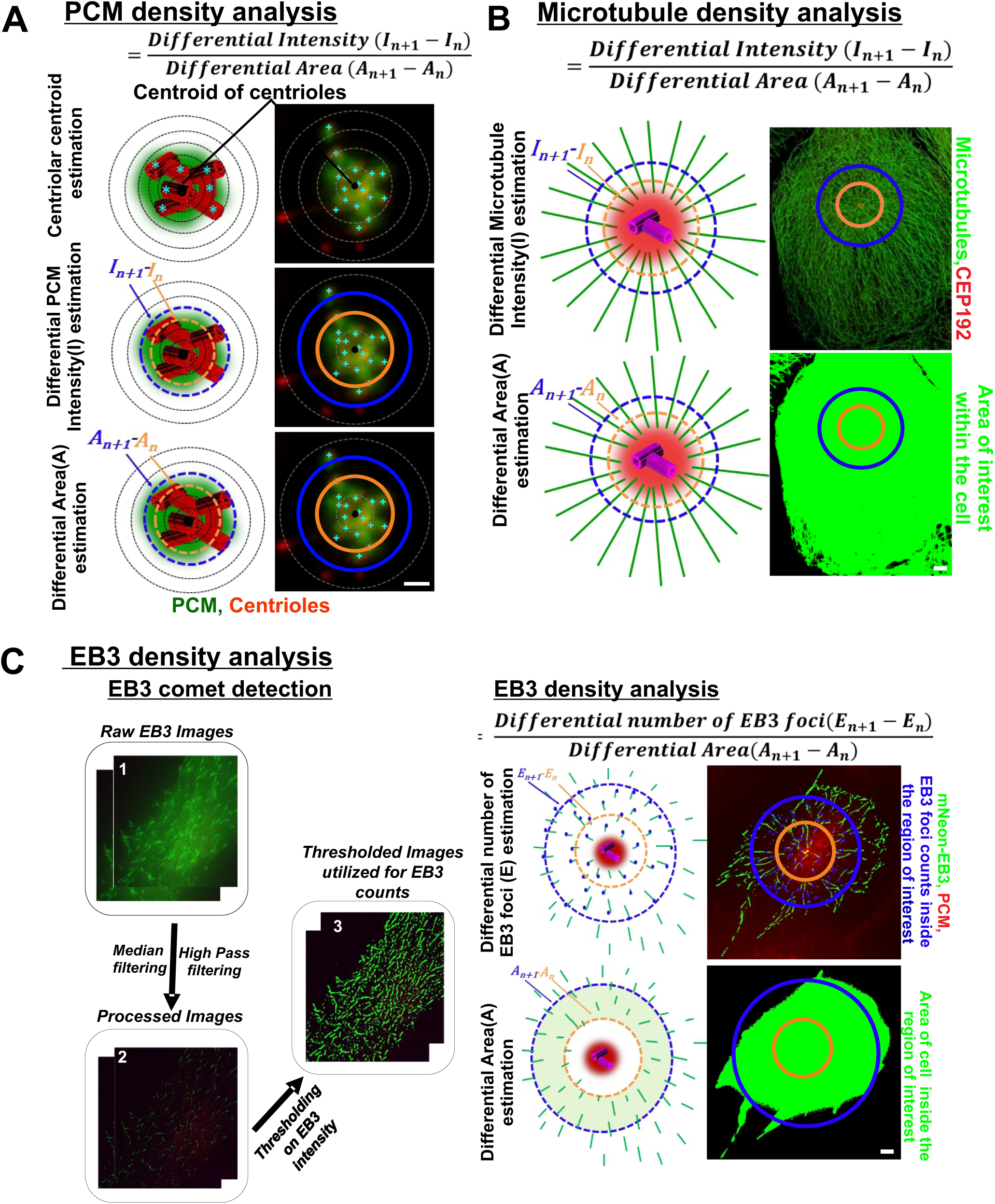
Algorithms to quantify PCM and microtubule density around centrioles. **(A)** PCM density analysis. Top left panel, schematic of centriolar centroid (black circle) estimation from centrioles (cyan foci). Top right panel, centrosomes in MDA-231 cells stained for centrioles (centrin, red) and γ-tubulin (green). Middle left panel, schematic for analysis of PCM intensity. Middle right panel, the region around the centroid of the centrioles is divided into concentric rings, and the differential intensity (In+1-In) of pericentriolar material is calculated around centrosomes (right) in MDA-231 cells. Bottom left panel, schematic for analysis of differential area (An+1-An). Bottom right panel, centrosomes (right) in MDA-231 cells stained for centrioles (centrin, red) and γ-tubulin (green). Scale bar represents 1 μm. **(B)** MT density analysis. Top left panel, schematic of analysis of differential MT intensity (In+1-In) for each ring. Top right panel, centrosomes in MDA-231 cells stained for centrioles (CEP192, red) and MTs (α-tubulin, green). Bottom left panel, schematic (left) for analysis of differential area (An+1-An) around centrosomes (right) in MDA-231 cells. Bottom right panel, the image of thresholded MTs. Scale bar represents 5 μm. **(C)** EB3 density analysis. Left, the pipeline for detecting EB3 foci. 1 original image, 2 processed image after applying the median filter and subtracting a blurred version of the same image, blurred using a box filter. Pixels on EB3 foci have high intensity and all other pixels have been made close to 0 intensity. 3 thresholded binary image. Connected components in this image are considered as EB3 foci. Right, schematic of analysis of EB3 density for each ring. Top left panel, schematic (left) for analysis of differential EB3 counts (En+1-En). Top right panel, centrosomes (right) in tet-induced mNeon-EB3 MCF10A cells stained for centrioles (centrin, green) and γ-tubulin (red). Bottom left panel, schematic for analysis of the differential area, Bottom right panel, the image of thresholded EB3 foci. Scale bar represents 5 μm.

Below we first describe the image processing algorithms for the analyses of centrosomal proteins, MTs, and EB3 comets. We then describe the semi-automatic graphical tool that encompasses both these image processing algorithms and the machine learning aided detection of centrosomes and cells.

#### PCM and MT Intensity Analyses

To quantify PCM and MT intensity in concentric rings at and around centrosomes, images were projected to their maxima and the centroid of the identified centrosomes was algorithmically computed. Next, the distance of each pixel from the centrosome centroid in two dimensions was calculated (Figure 2A top panel). For each radius from inner r1 to rn where n is the user-defined number of radii used in the analysis, the tool computes the total fluorescence intensity (In) of the pixels between the concentric radii. The difference in total intensity (In+1 – In) between the n-th and (n+1)-th regions provides the fluorescence intensity in the n-th concentric ring (Figure 2A middle panel, 2B top panel; n-th: orange and (n+1)-th: blue concentric ring). To correct for the total area of each ring the intensity is normalized to the number of pixels within each ring.

Only pixels that fall within the cell boundary are quantified. To define the cell boundary for the MT intensity computation, a low threshold was used on the α −tubulin fluorescence channel; this was motivated by the fact that tubulin fluorescence extends throughout the cell area (Figure 2B bottom panel). In case such a channel is not available, one can also use the cell body estimated by the cell segmentation algorithm as a rough estimate. For the PCM density computation, smaller radii (in steps of 0.13 μm) and a smaller total diameter was used so that the analysis region was always inside the cell (cells with centrosomes very close to edge of the cell/ edge of the image were not analyzed).Therefore, we did not need to account for the cell area (Figure 2A). In each case, once the pixels within the cell were defined (i.e. cellular area) then, for each radius from inner r1 to rn the tool computes the differential area (An+1 – An) between the n-th and (n+1)-th regions. Dividing the differential intensity by the differential cell area provided the density (Figure 2B). Altogether, this generated an analysis routine to measure the spatial distribution of MTs and PCM fluorescence per unit area.

#### EB3 Foci Analysis

To quantify the distribution of MT nucleation and growth from centrosomes, we designed a similar algorithm to measure the distribution of EB3 foci on images from fixed cells (Figure 2C). The algorithm detects EB3 comets by utilizing a median filtering^2^ step to remove noise in the form of stray pixels with high fluorescence intensity. Then, to identify pixels with greater intensity than their neighbors, *high-pass filtering* was performed where a blurred version of the image (obtained by convolving^3^ the image with the box filter^4^) was subtracted from the image. Thresholding this differential image provided a binary image that identified pixels corresponding to EB3 comets. Finally, each connected component^5^ in this binary image was considered an EB3 comet (Figure 2C).

Analyzing the distribution of EB3 comets in concentric rings as described above requires additional considerations since EB3 comets can straddle multiple rings. For each EB3 comet, we identified the pixel on the comet farthest from the centroid of the centrosomes. Then, for each radius (rn), we counted the EB3 comets for which this pixel fell within a distance of rn from the centroid of centrosomes. The differential EB3 counts were then computed in the n^th^ ring as the difference in comet counts (En – En+1) and divided by the differential cell area to obtain the EB3 density (Figure 2C right panel).

#### A semi-automatic graphical tool for MT, PCM and EB3 analyses

We encapsulated the centrosome detection, cell segmentation, and MT/PCM and EB3 analysis algorithms described above into a single tool with a graphical user interface (Figure 3A). This graphical user interface does not include the “training” of the machine learning-based centrosome detector, which is trained separately and used for all subsequent analyses.

**Figure 3:**
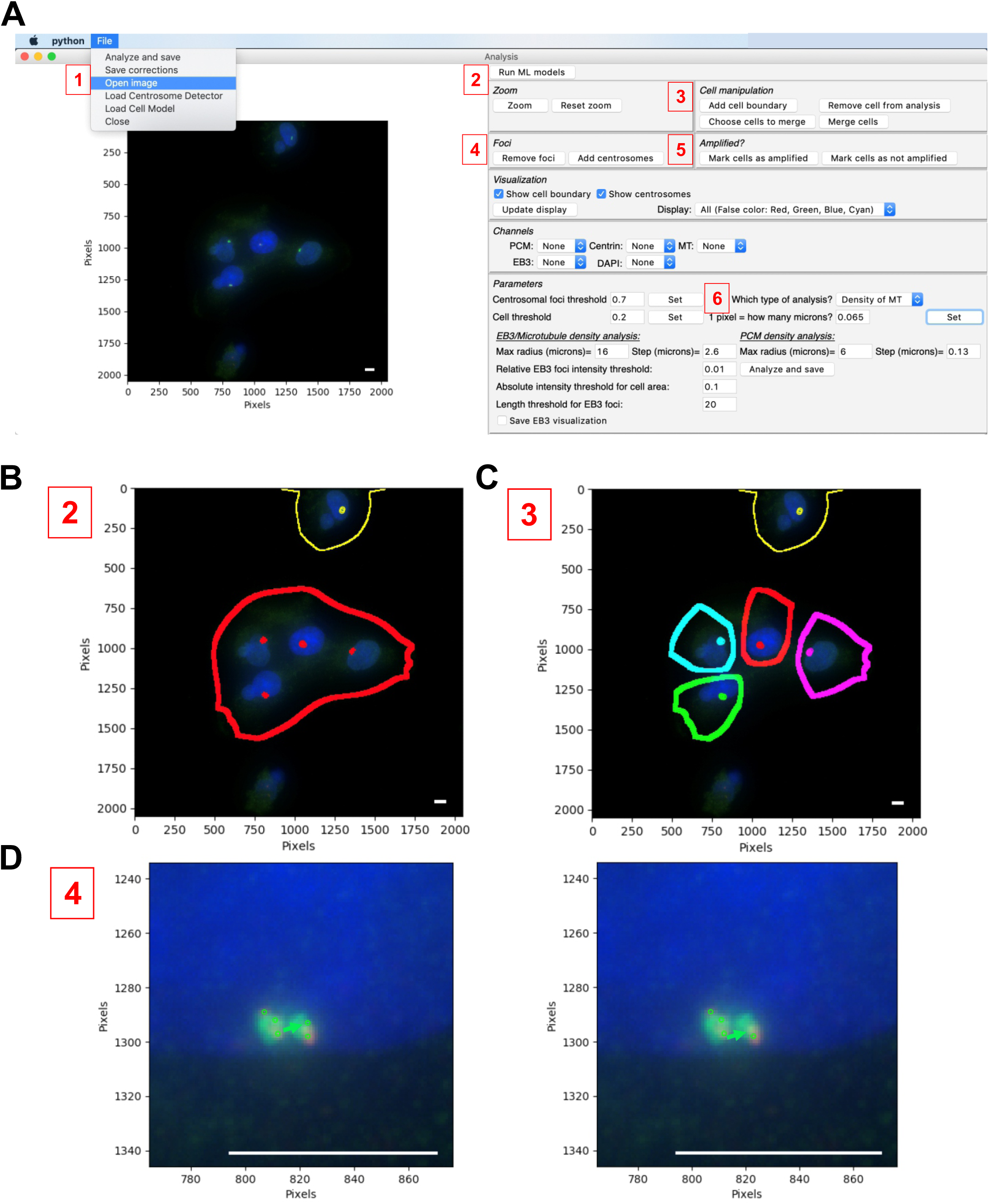
A semi-automatic graphical tool for performing PCM and MT density analysis **(A)** Screenshot of graphical interface that encapsulates machine learning aided detection of centrosomes with image processing algorithms to analyze the spatial distribution of MT organization. The steps to perform the PCM and MT density analysis are shown in red boxes. Scale bar represents 10 μm. **(B)** Screenshot of an image in the graphical interface with centrosomes (red, cyan, and yellow foci) and cell boundaries (red, cyan, and yellow boundaries) detected. **(C)** Screenshot of the image in the graphical interface showing edited cell boundaries (red, green, magenta, white, cyan, and yellow boundaries). Scale bar represents 10 μm. **(D)** Left panel, screenshot of a spurious detection (green arrow) as a centrosome. Right panel, screenshot after removal of the spurious detection (green arrow) in the graphical interface. Scale bar represents 10 μm.

To ensure performance, this is a semi-automatic tool that allows for manual intervention. In particular, the user can choose thresholds for centrosome detection and cell segmentation and visualize the machine’s outputs. The user can correct these outputs by removing spurious foci, including centrosomes that the algorithm missed, adding cell boundaries that were not detected and merging cells that have been mistakenly marked as two separate cells. The tool also allows the user to control various aspects of the analysis itself, such as the radii for spatial analysis, the thresholds for identifying EB3 comets, and the thresholds for determining the cell area. Finally, this tool can be easily generalized to analyze the distribution of any protein around centrosomes or any other user-identified foci.

With this tool, analysis is performed with the following steps. First, the image, centrosome detection model and cell segmentation model are loaded (Figure 3A; step 1). Second, the centrosome detection and cell segmentation models run (run Machine Learning (ML) models; Figure 3A; step 2). This will automatically detect centrosomes and cell boundaries and present these as an overlay on the original image (Figure 3B). In case of an error in the cell segmentation, the segmentation can be corrected by adding additional cell boundaries or merging wrongly segmented cells. One can also remove cells from further analysis (Figure 3C; step 3). The tool also allows for the removal of spurious detections or the inclusion of missed centrosomes (Figure 3D; step 4). The tool automatically characterizes cells as being centrosome-amplified based on the number of detected centrosomes (cells with greater than two centrosomes), with amplified cells highlighted by a thick outline. This automated labeling can also be corrected if needed (Figure 3A; step 5). Finally, once the centrosomes have been finalized, the type of analysis is chosen (Figure 3A; step 6). The analysis is automatically performed and the results are saved in a tabular format.

The tool also allows for corrections (i.e., removal of spurious detections and identification of missed centrosomes) to be saved for future training of the centrosome detection model. The training can be performed using a separate command-line utility that is packaged with the tool.

In summary, we have generated a new tool with a graphical interface that allows users to analyze the spatial distribution of centrosomal proteins and MTs around centrosomes (https://bharath272.github.io/centrosome-analysis/). The tool performs automated analyses but also allows for user interventions and training of the underlying machine learning models.

### Semi-automatic machine learning algorithm detects PCM defects in breast cancer cells

To test whether PCM levels are elevated in breast cancer cells with amplified centrosomes, the above-described algorithm was used to quantify the levels and distribution of PCM proteins in normal and breast cancer cells (MCF10A and MDA-231, respectively).

Analyses were first performed using manual annotation of centrosomes. The levels and distribution of PCM proteins (γ-tubulin and Pericentrin) were quantified at centrosomes with normal or amplified numbers. Approximately 23% of MDA-231 breast cancer cells (MDA-231) have CA while only 5% of normal-like breast cells (MCF10A) have CA (Ganapathi Sankaran et al., 2019). The relative intensities of *γ*-tubulin and Pericentrin were quantified per unit area in normal compared to amplified centrosomes (Figure 4). Centrosomal *γ*-tubulin was elevated by approximately 40% at amplified centrosomes compared to non-amplified centrosomes in both MCF10A and MDA-231 cells (Figures 4A and 4B). The relative *γ*-tubulin intensity diminishes by at least 70% outside the pericentriolar space (between 1.0 to 2.0 μm) similar to the non-amplified centrosomes (Figure 4B). This suggests that while the PCM protein levels are elevated at the core centrosome, *γ*-tubulin remains constrained to the pericentriolar space of the amplified centrosomes. Consistent with the increase in *γ*-tubulin intensity, Pericentrin protein is also elevated in breast cancer cells with amplified centrosomes (Figure 4C and 4D). However, unlike *γ*-tubulin, the Pericentrin protein intensity at the amplified centrosomes does not diminish as rapidly outside the core pericentriolar space when compared to non-amplified centrosomes.

**Figure 4:**
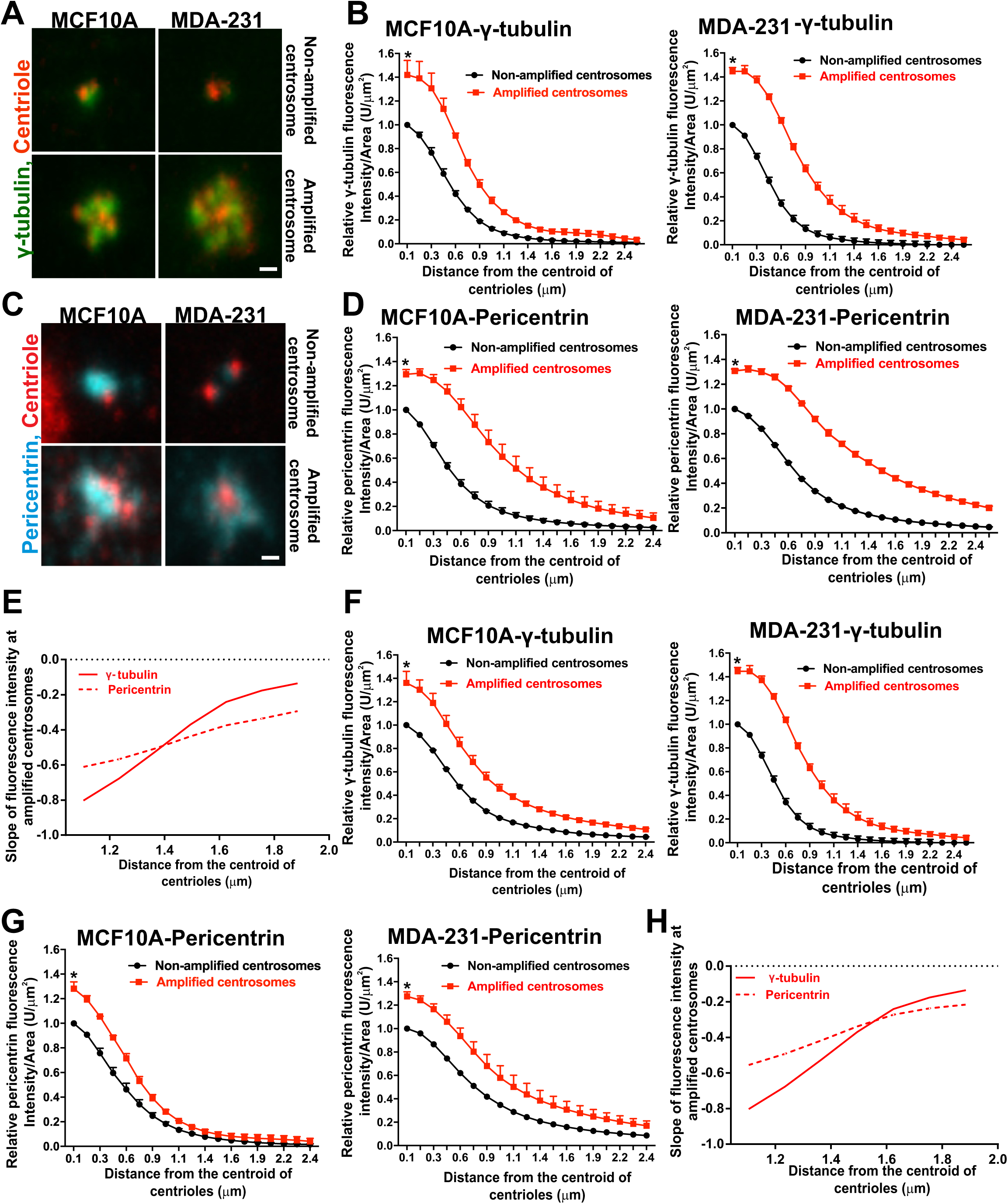
Centrosome amplified breast cancer cells exhibit PCM defects. **(A)** Left panel, non-amplified and amplified centrosomes in MCF10A cells stained for centrioles (centrin, red) and γ-tubulin (green). Right panel, non-amplified and amplified centrosomes in MDA-231 cells stained for centrioles (centrin, red) and γ-tubulin (green) Scale bars represent 1 μm. **(B)** Left panel, γ-tubulin fluorescence density in amplified (red) cells relative to non-amplified (black) MCF10A cells (p-value: 1.9*10^-6^). Right panel, γ-tubulin fluorescence density in amplified (red) cells relative to non-amplified (black) MDA-231 cells (p-value: 1.9*10^-6^). **(C)** Left panel, non-amplified and amplified centrosomes in MCF10A cells stained for centrioles (centrin, red) and Pericentrin (cyan). Right panel, non-amplified and amplified centrosomes in MDA-231 cells stained for centrioles (centrin, red) and Pericentrin (cyan). Scale bars represent 1 μm. **(D)** Left panel, Pericentrin fluorescence intensity in amplified (red) cells relative to non-amplified (black) MCF10A cells (p-value: 3.8*10^-6^). Right panel, Pericentrin fluorescence intensity in amplified (red) cells relative to non-amplified (black) MDA-231 cells (p-value: 1.9*10^-6^). **(E)** The slope or first derivative of fluorescence intensities of γ-tubulin and Pericentrin at amplified centrosomes outside the core pericentriolar space (between 1.0 μm to 2.0 μm). **(F)** Left panel, γ-tubulin fluorescence intensity in amplified (red) cells relative to non-amplified (black) MCF10A cells obtained using machine annotation (p-value: 3.8*10^-6^). Right panel, γ-tubulin fluorescence intensity in amplified (red) cells relative to non-amplified (black) MDA-231 cells obtained using machine annotation (p-value: 3.8*10^-6^). **(G)** Left panel, Pericentrin fluorescence intensity in amplified (red) cells relative to non-amplified (black) MCF10A cells obtained using machine annotation (p-value: 3.8*10^-6^). Right panel, Pericentrin fluorescence intensity in amplified (red) cells relative to non-amplified (black) MDA-231 cells obtained using machine annotation (p-value: 3.8*10^-6^). **(H)** The slope or first derivative of fluorescence intensities of γ-tubulin and Pericentrin at amplified centrosomes outside the core pericentriolar space (between 1.0 μm to 2.0 μm). Mean±SEM. Wilcoxon test.

To quantify the differences in how *γ*-tubulin and Pericentrin diminish outside the pericentriolar cloud, we estimated the first derivative or slope of *γ*-tubulin and Pericentrin fluorescence intensities outside the core pericentriolar space (Figure 4E). The slope of *γ*-tubulin intensity at amplified centrosomes is more negative in comparison to the slope of Pericentrin intensity at amplified centrosomes. *γ*-tubulin fluorescence intensity diminishes quickly outside the core pericentriolar space at amplified centrosomes in comparison to Pericentrin fluorescence intensity. Moreover, this suggests that the increase in centrioles at amplified centrosomes modulates the organization of specific pericentriolar proteins.

We next utilized the semi-automatic machine learning algorithm to quantify the pericentriolar defects at amplified centrosomes that were described using manual annotation and quantification above. The semi-automatic graphical tool annotated cells with greater than two centrosomes as amplified. However, if the cells were mis-annotated as non-amplified, then the annotations were manually edited to amplified centrosomes. The tool corroborated the results from the manual quantification: higher γ-tubulin levels relative to non-amplified centrosomes were reported by the semi-automated analysis (Figure 4B and 4F). While the *γ*-tubulin intensity is elevated, the distribution remains encompassed within the pericentriolar space (60% of total fluorescence intensity) of the amplified centrosomes. Pericentrin is also elevated in breast cancer cells with CA (Figure 4D and 4G). Furthermore, Pericentrin expands the pericentriolar space of amplified centrosomes (Figure 4H). These results show that the semi-automatic machine learning algorithm detects the same centrosome organization defects found at amplified centrosomes as found by the more laborious manual annotation and quantification.

### Increased MTs from amplified centrosomes

The increase in *γ*-tubulin at amplified centrosomes suggests that MTs may also be elevated around centrosomes. To test whether cells with amplified centrosomes have increased MTs, we measured the MT fluorescence intensity per unit area using the algorithm. MT intensity is 30% greater in both MCF10A and MDA-231 cells with amplified centrosomes compared to cells without CA (Figure 5A and 5B). Overall, these data suggest that centrosome amplified breast cancer cells have an increase in centrosome-associated *γ*-tubulin that promotes MT nucleation to increase the MT density.

**Figure 5:**
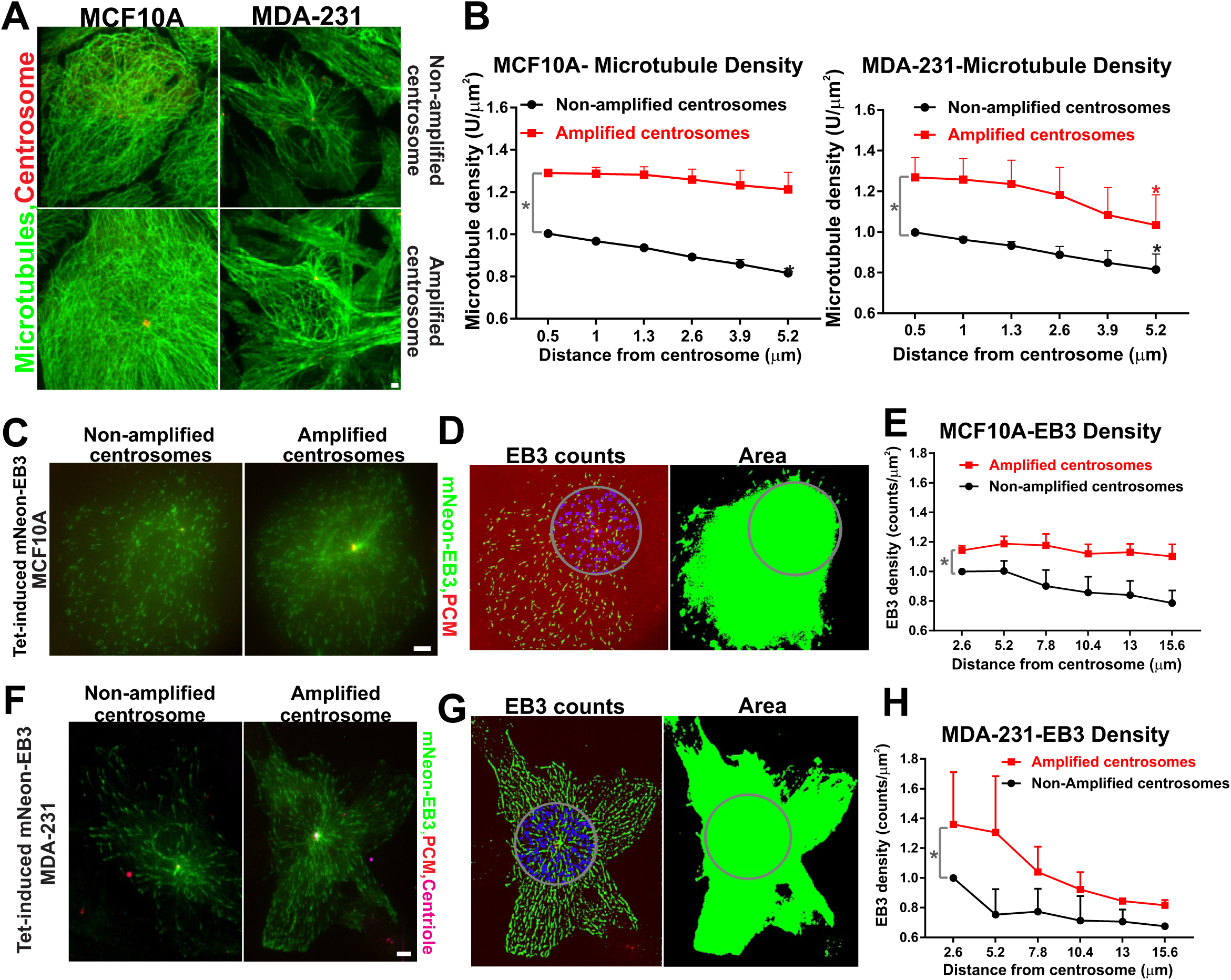
Centrosome amplified breast cancer cells exhibit increased MT density. **(A)** Left panel, non-amplified and amplified centrosomes in MCF10A cells stained for centrioles (CEP192, red) and MTs (α-tubulin, green). Right panel, non-amplified and amplified centrosomes in MDA-231 cells stained for centrioles (CEP192, red) and MTs (α-tubulin, green). Scale bars represent 10 μm. **(B)** Left panel, MT fluorescence intensity in amplified (red) MCF10A cells relative to non-amplified (black) cells (p-value; gray asterisk: 4*10^-6^, black asterisk: 3.4*10^-6^). Right panel, MT fluorescence intensity in amplified (red) MDA-231 cells relative to non-amplified (black) cells (p-value; gray asterisk: 8*10^-6^, black asterisk: 2*10^-2^, red asterisk; 9.3*10^-3^). **(C)** Non-amplified and amplified centrosomes in Tet-induced mNeon-EB3 MCF10A cells stained for centrioles (centrin, green) and γ-tubulin (red). Scale bars represent 5 μm. **(D)** Processed images of non-amplified Tet-induced mNeon-EB3 MCF10A cells utilized for the measurement of EB3 counts and area. **(E)** EB3 foci counts per unit area in non-amplified (black) and amplified (red) MCF10A cells (p-value: 3.1*10^-2^). **(F)** Non-amplified and amplified centrosomes in Tet-induced mNeon-EB3 MDA-231 cells stained for centrioles (centrin, green) and γ-tubulin (red). Scale bars represent 5 μm. **(G)** Processed images of amplified Tet-induced mNeon-EB3 MDA-231 cells utilized for the measurement of EB3 counts and area. **(H)** EB3 counts per unit area in amplified (red) MDA-231 cells relative to non-amplified (black) cells (p-value: 3.1*10^-2^). Mean±SEM. Wilcoxon test and Students t-test (paired).

To test whether amplified centrosomes increase the number of growing MT ends, we created stably expressing tetracycline-inducible mNeon-EB3 MCF10A and MDA-231 cell lines. Tetracycline was added for 48 hours to express mNeon-EB3 and the number of EB3 comets per unit area was measured using the EB3 density feature in the graphical interface (Figure 5C-H). A 20% and 40% increase in the EB3 density was observed at amplified centrosomes relative to the non-amplified centrosomes in MCF10A and MDA-231 cells, respectively. This suggests that amplified centrosomes nucleate more MT ends compared to non-amplified centrosomes. Furthermore, the density of EB3 foci is elevated closer to centrosomes in comparison to the cell periphery in both MCF10A and MDA-231 cells (Figure 5E and 5H). This is consistent with the elevated MT density that is observed closer to centrosomes relative to the cell periphery (Figure 5B). Taken together, the data suggest that the MT nucleation is enriched at amplified centrosomes in comparison to the non-amplified centrosomes of MCF10A and MDA-231 cells.

## DISCUSSION

### A semi-automatic graphical tool for centrosome detection and analysis of MT organization

We designed a semi-automatic graphical tool that can significantly speed and aid in removing bias in the analysis of centrosomes and MTs as it removes the need for the user to manually identify centrosomes or EB3 comets. Because of limitations in the amount of training the machine learning algorithm can realistically incorporate, we found that user correction of the algorithm’s predictions was crucial to ensure the correctness of the analysis. These corrections can be used to further train the underlying model, allowing the model to improve with use. However, in our experiments, we did not perform such retraining so as to keep the model fixed during analysis. Furthermore, retraining provides flexibility to the tool, allowing it to be used for different experiments. Training scripts and models are publicly available to the research community (https://bharath272.github.io/centrosome-analysis/).

While correcting the output from the algorithm, we found several common modes of error. First, unlike internet images where machine-learning technologies are routinely used, the dynamic range of fluorescence images can vary significantly, and naïve training of convolutional networks does not generalize across such variations. We addressed this by normalizing each image independently to have zero mean and unit variance. Second, we found that while the algorithm often correctly identified the rough location of the centrosomes, it often did not correctly estimate their number. We thus had to correct its predictions as to which cells exhibited CA. Accurate counting has not been addressed in the computer vision literature and deserves further attention (Chattopadhyay et al., 2017). Finally, we found that the algorithm often merged nearby cells or estimated their boundary incorrectly. These issues reveal challenges that must be addressed by future computer vision research. Nevertheless, our results show that existing techniques can provide a significant degree of automation and improvement to the limited efficiency and biases associated with manual annotation.

### PCM proteins and their organization around amplified centrosomes in breast cancer

Taxol/Paclitaxel, Taxotere, and Cabazitaxel are MT stabilizing drugs that have been used in chemotherapy for the treatment of cancers (Ho and Mackey, 2014; Pazdur et al., 1993; Schiff and Horwitz, 2006). Taxol and other MT stabilizing drugs lead to mitotic arrest. However, the proportion of cells lost by mitotic arrest does not explain the efficacy of taxol as a chemotherapy agent. The alternate hypothesis that taxol affects interphase breast cancer cells has not been well studied (Weaver, 2014). Specifically, it is unclear whether these MT stabilizing drugs differentially impacts underlying differences in the interphase centrosomes and MTs of normal compared to transformed breast cells (Weaver, 2014). To understand the differences in interphase centrosomes and MTs of breast cancer cells, we investigated the levels and the distribution of centrosome PCM proteins and MTs using our semi-automatic machine learning-aided approach. We find that amplified centrosomes in interphase breast cancer cells not only have elevated *γ*-tubulin and Pericentrin but Pericentrin also has an expanded distribution. This suggests that there are more PCM sites that promote MT nucleation and MTs that are organized around amplified centrosomes. This may disrupt MT-related processes like cellular motility, signaling, and MT-dependent transport (Caviston and Holzbaur, 2006; Desai and Mitchison, 1997; Hyman and Karsenti, 1996; Inoue’ and Salmon, 1995). Furthermore, expansion of the PCM may interfere with the MT-motor-dependent trafficking (Galati et al., 2018; Nanjundappa et al., 2019). This altered MT-motor-dependent trafficking may affect drug transport and sensitivity to MT stabilizing drugs. Taken together, the centrosome amplified population with increased MT organization may have a role in taxol toxicity/chemo-resistance.

### The interphase MT network is altered in centrosome amplified breast cancer cells

In addition to the increased PCM, the density of MTs is elevated in MCF10A and MDA-231 cells with amplified centrosomes compared to non-amplified centrosomes. The increased MT organization could be attributed to cell cycle differences since S phase cells have elevated microtubule density relative to G1 phase cells (Salaycik, 2005). However, the pool of non-amplified centrosomes includes interphase cells with both one (likely G1 phase) or two (likely S phase) centrosomes. Hence, cell cycle differences likely do not explain the increase in MT organization observed in the centrosome amplified cell population. The increase in microtubule nucleation from additional PCM sites is a more likely explanation for the increase in MT density that is observed with centrosome amplification. Finally, whether the cytoplasmic α/β-tubulin concentrations are elevated in these cells to promote increased MT assembly is not known. Nevertheless, our studies suggest that amplified centrosomes promote an increased MT network in interphase breast cancer cells.

We investigated whether the number of growing MTs, as judged by EB3 density, is increased in breast cancer cells with amplified centrosomes. Indeed, the EB3 density is elevated in centrosome amplified breast cancer cells relative to the cells without centrosome amplification (Figure 5C-D). Furthermore, EB3 density decreases towards the cell periphery of centrosome amplified MDA-231 cells relative to centrosome amplified MCF10A cells. This is consistent with the observation that MT density decreases towards the cell periphery of centrosome amplified MDA-231 cells. In contrast, MT density in MCF10A cells with amplified centrosomes is greater at the cell periphery (Figure 5B). The increased MT density from MCF10A amplified centrosomes at the cell periphery might be explained either by the presence of longer and more stable MTs or by increased nucleation and branching of the MT network (Goshima et al., 2008; Ishihara et al., 2014; Petry et al., 2013). It remains unclear why MCF10A and MDA-231 cells exhibit these unique profiles in MT growth and density. An increase in MT density and EB3 comets around amplified centrosomes reflect differences in MTs between non-amplified and amplified interphase breast cancer cells. Such altered cytoskeletal architecture could lead to changes to intracellular trafficking and disrupt important cellular processes including ciliogenesis, cell migration and cell polarization (Bouchet and Akhmanova, 2017; Caviston and Holzbaur, 2006; Siegrist and Doe, 2007).

In summary, we built a semi-automated tool with a graphical interface that enables quantitative measurements of centrosome aberrations. Using this approach, we detect centrosome aberrations in cancer cells and show an increase in MT nucleation that promotes changes to MT density and the spatial distribution of MTs around centrosomes. This not only highlights the use of the machine learning-based approaches for the facile detection of centrosomal aberrations but also reveals how it can be globally applied towards quantitative cell biological problems.

## MATERIALS AND METHODS

### Cell Culture

Breast cancer cell lines MCF10A and MDA-MB-231 (MDA-231) were obtained from the University of Colorado Cancer Center Tissue Culture Core. Mammalian tissue culture lines were all grown at 37°C with 5% CO_2_. MCF10A cells were received at passage 51 and were grown in DMEM/F12 (Invitrogen #11330-032), 5% Horse Serum (Invitrogen #16050-122), 20 ng ml^-1^ EGF (Invitrogen #PHG0311), 0.5 mg ml^-1^ Hydrocortisone (Sigma #H-0888), 100 ng ml^-1^ Cholera toxin (Sigma #C-8052), 10 μg ml^-1^ Insulin (Sigma #I-1882) and 1% Pen/Strep (Invitrogen #15070-063). MDA-MB-231 cells were received at passage 15 were grown in DMEM (Invitrogen #11965-092), Pen/Strep (Invitrogen #15070-063) and 10% FBS (FBS; Gemini Biosciences). Cell lines were authenticated at the sources and tested negative for mycoplasma using the MycoAlert mycoplasma detection kit through the University of Colorado Cancer Center Tissue Culture Core. Cells were passaged and sub-cultured using Trypsin (Invitrogen #150901-046) when cultures reached 60-80% confluency.

### Generation of Tetracycline Inducible mNeon-EB3-MCF10A and mNeon-EB3-MDA-231 cells

The generation of the tetracycline-inducible mNeon-EB3 construct is described below. The EB3-mNeon (C-terminal fusion) fragment was obtained through PCR with Phusion DNA polymerase of a pre-existing plasmid using primers that have Nhe1 and Xma1 sites appended to them. This was cloned into the tetracycline-inducible construct pcw57.1 using the enzymes Nhe1 and Age1.

Lentivirus harboring tetracycline-inducible mNeon-EB3-MCF10A and MDA-231 were made by the transfection of 293FT cells. 293FT cells were plated in 6 cm dishes and allowed to reach 50%-70% confluency. Cells were then transfected with tetracycline-inducible EB3-mNeon constructs, and second-generation lentivirus packaging plasmids (pMD2.G and psPAX2) using Lipofectamine 2000 (Life Technologies # 11668019). 293FT media containing virus was harvested and MDA-231 cells were infected for 24-48 hours in the presence of 10 μg ml^−1^ (26.7 μM) polybrene. After a 24-hour recovery, transduced cells were selected with puromycin at 2 μg ml^−1^ (4.24 μM) and were flow-sorted to isolate and plate single cells into 96 well plates. Such clones were cultured in 50% filtered conditioned media with 50% fresh media. Tetracycline-inducible mNeon-EB3-MCF10A and MDA-231 cells were induced with tetracycline (Invitrogen #550205) at 2.5 μg ml^−1^ (5.63 μM).

### Transfections

MCF10A and MDA-231 cells at 50-80% confluence were transfected using Lipofectamine 2000 (Invitrogen # 11668019). Plasmid DNA and Plus reagent (Invitrogen # 11514015) were mixed at 1:1 and incubated for 5 minutes. This mixture was then combined with Lipofectamine at a1:3 ratio. Complexes were diluted in Opti-MEM (Invitrogen # 31985062). After a 4-hour incubation, the complexes were removed and the transfected cells were supplied with fresh media.

### Immunofluorescence

12 mm diameter coverslips were acid-washed and heated to 50°C in 100mM HCl for 16 hours. This was followed by washes with water, 50%, 70%, and 95% ethanol for 30 minutes each. Coverslips were coated with Type-1 collagen (Sigma # C9791), air-dried for 20 minutes in the laminar hood and exposed to UV light for cross-linking of collagen for 20 minutes. Cells were cultured on collagen-coated coverslips to 55-70% confluence. For centrosome immunofluorescence, cells were fixed with 100% methanol at −20°C for 8 minutes. Fixed cells were washed with PBS/Mg (1x PBS, 1mM MgCl2), and then blocked with Knudsen Buffer (1x PBS, 0.5%BSA, 0.5% NP-40, 1mM MgCl2, 1mM NaN3) for 1 hour. Cells were incubated overnight with primary antibodies diluted in Knudsen Buffer at 4°C. Coverslips were washed with PBS three times in 5-minute intervals. Secondary antibodies and Hoechst 33258 (10μg ml^−1^, Sigma #B2261) were diluted in Knudsen buffer and incubated for 1 hour at room temperature. Coverslips were mounted using Citifluor (Ted Pella) and sealed with clear nail polish. Antibodies used for immunofluorescence are α-centrin (1:2,000; 20H5; Abcam), *α*-*γ*-tubulin (1:1000; DQ-19; Sigma), α-α-tubulin (1:500; DM1A; Sigma). Alexa-fluor secondary antibodies were diluted to 1:1,000 for all experiments (Molecular Probes).

### Microscopy

The fluorescence imaging utilized for Figures is identical to those described in Dahl et al, 2015. Briefly, images were acquired using a Nikon TiE (Nikon Instruments, Inc.) inverted microscope stand equipped with a 100X PlanApo DIC, NA 1.4 objective. Images were captured using an Andor iXon EMCCD 888E camera or an Andor Xyla 4.2 CMOS camera (Andor Technologies). Images in Figure 4A-B were acquired using a Swept Field Confocal system (Prairie Technologies / Nikon Instruments) on a Nikon Ti inverted microscope stand equipped with a 100X Plan Apo λ, NA 1.45 objective. Images were captured with an Andor Clara CCD camera (Andor Technologies).

Nikon NIS Elements imaging software was used for image acquisition. Image acquisition times were constant within a given experiment and ranged from 50 to 400 msec, depending on the experiment. All images were acquired at approximately 25°C. Images presented in most of the figures are maximum-intensity projections of the complete *z*-stacks. Exceptions include certain mitotic images that are constructed from selected z-planes to clearly distinguish kinetochores and lagging chromosomes.

### Centriole and centrosome number counts

Cells were scored as amplified, non-amplified and under duplicated based on centrin and *γ*-tubulin staining (Dahl et al., 2015). Cells with greater than two *γ*-tubulin and four centrin foci were scored as amplified centrosomes. Non-amplified centrosomes have both one or two *γ*-tubulin and two or four centrin foci.

### Computational tools

The semi-automatic graphical user interface was created as a standalone python program. The training script was a separate program packaged with the graphical interface. The version of python used was 3.6, and it was used in conjunction with Anaconda as a package manager. The graphical user interface accepts TIFF files. The size that a pixel denotes can be input through the interface. The graphical user interface was created using the Tkinter Python library. The underlying machine learning models were built using the PyTorch library; we used version 0.4.1. The code also relies on the numpy library for matrix computations, the scipy library for a variety of signal processing routines, the matplotlib library for plotting and the tifffile library for reading and writing tiff images. The training annotations were collected using a separate stand-alone program with a graphical user interface.

The centrosome detection model was trained using stochastic gradient descent with momentum. The parameters of the training procedure were as follows: learning rate: 0.01, momentum: 0.9, weight decay: 0.0001, the total number of epochs: 100, batch size: 1. Because pixels on centrosomes are very few in number, the learning problem was found to be extremely imbalanced. To deal with this imbalance, the training loss on centrosomal pixels was increased by a factor of 1000. This is mathematically equivalent to statistically sampling centrosomal pixels 1000 times more often during the stochastic gradient descent procedure. Pixels within 5 pixels of a centrosome were ignored during training to avoid penalizing the algorithm for small deviations.

The cell segmentation model was also trained with stochastic gradient descent with momentum. The parameters of the training procedure were as follows: learning rate: 0.01, momentum: 0.9, weight decay: 0.0001, the total number of epochs: 100, batch size: 1. The tolerance for the random walk segmenter was set to 0.01. The code is available here: https://bharath272.github.io/centrosome-analysis/

Also included with the code is a training script that can be used to train a new model based on the saved corrections from the graphical tool. This training script can be run from the terminal as follows:

python train_foci_detector.py --trainfiles trainingset.csv --modelfile <u>foci_model.pt</u> Here, trainingset.csv must be a CSV (comma-separated values) file formatted as follows. Each row of this file consists of the path to the image (a TIFF file), and the path to the corresponding corrected annotation (a JSON file, saved from the graphical tool using the “Save corrections” feature), separated by a comma. Such a CSV can be created using most spreadsheet software including Microsoft Excel. Once run, the training script will save the resulting trained foci detector in the file <u>foci_model.pt</u>.

### Statistics and biological replicates

All center values represent means and error bars represent the standard error of the mean. All the experiments were performed using at least three independent biological replicates. The number of cells used in each immunofluorescence experiment is as follows: 3A and 3E: 40 cells per condition/80 cells per experiment. 3C and 3F:40 cells per condition/80 cells per experiment. 4A: 30 cells per condition/ 60 cells experiment. 4B: 40 cells per experiment. 4C: 40 cells per experiment. Fischer’s exact test, Student’s two-tailed t-test, Mann-Whitney U-test, and Wilcoxon tests were used to assess statistical significance between means. Fischer’s test was utilized to examine the significance of contingency when data were classified into two or more categories. Student’s two-tailed unpaired t-test was used to examine significance between two normal distributions (equal variance assumed). Normality tests were performed both on the raw data and meta-data extracted from the replicates of raw data. Shapiro-Wilk normality test and D’Agostino-Pearson omnibus normality test was utilized to examine the normality of data. Shapiro-Wilk normality test was used when the number of samples was less than eight. When the number of samples was greater than eight, the D’Agostino-Pearson omnibus normality test was used. Mann-Whitney u-test was utilized to examine the significance of non-normal unpaired distributions. Wilcoxon test was used utilized to examine the significance of non-normal paired distributions. Results were considered statistically significant with p-values less than 0.05. P-values were denoted in the figure legends.

## ACKNOWLEDGMENTS

The authors would like to thank Alexander J Stemm-Wolf for helpful discussions and detailed feedback on the manuscript. We thank Drs. Heide Ford for viral constructs, Jeffery Moore for the EB3-mNeon construct, Andrew Holland for α-CEP192 antibodies, and UC Tissue culture core for the cell lines. C.G.P. is supported by ACS (RSG-16-157-01-CCG), NIGMS (GM099820) and the Boettcher Foundation.

## DECLARATIONS

The authors declare no competing issues.

## AUTHOR CONTRIBUTIONS

DGS designed, performed experiments and wrote the manuscript. BH designed the algorithms, the machine learning models and the graphical interface. BH and CGP supervised the project and wrote the manuscript.

---

**Figure S1:**
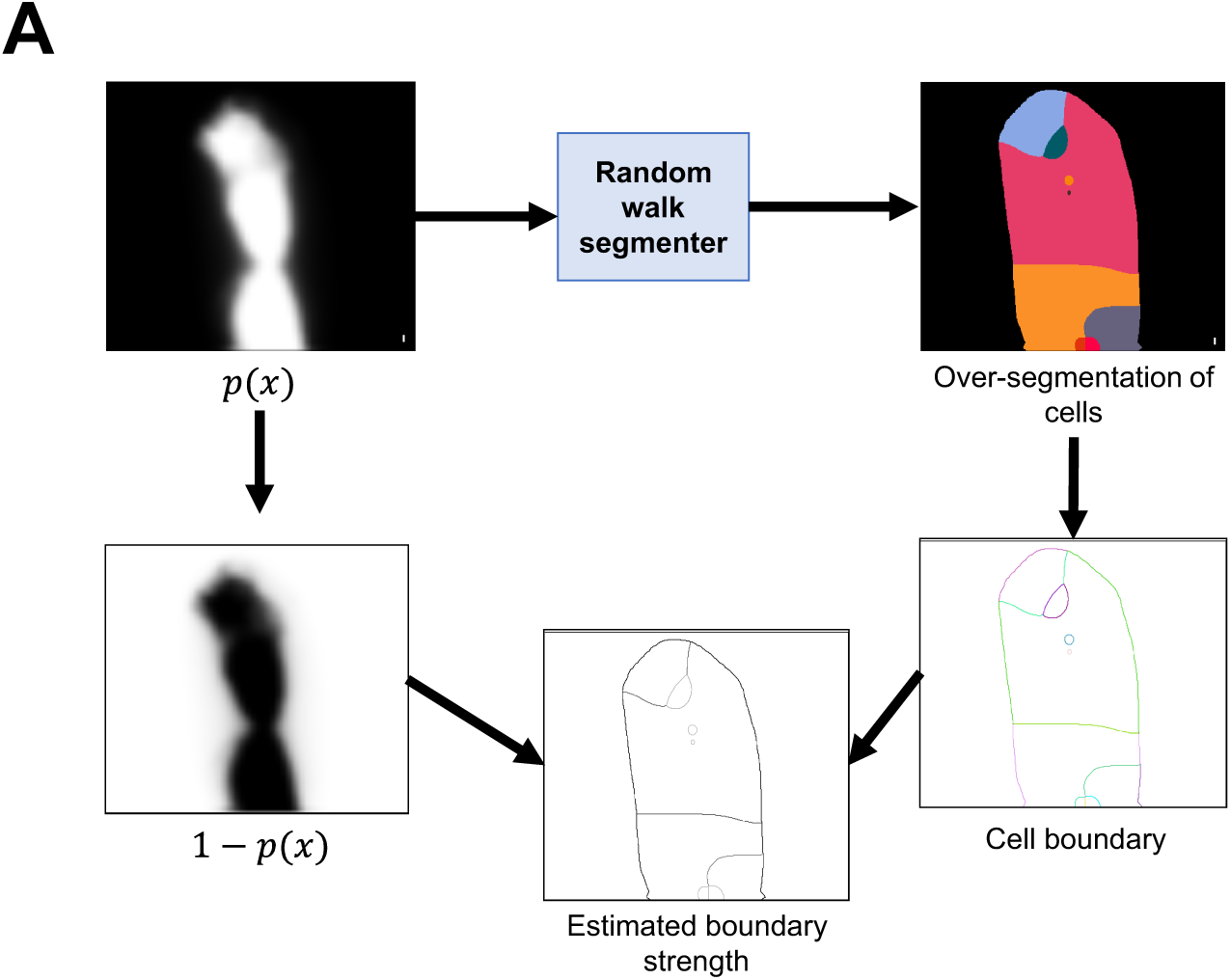
Pipeline for estimating boundary strength based on the convolutional network output, p(x). The random walk segmenter over-segments cells (top right). We identify the boundary between the predicted cells (bottom left) and estimate the strength of the boundary (bottom) using 1-p(x) (bottom right).

## Footnotes

1 **Convolutional networks**: Convolutional networks are machine learning models comprised of sequences of convolution operations interspersed with subsampling operations. The filters of these convolutions are automatically estimated by the learning algorithm based on a training dataset consisting of pairs of images and the desired outputs.

2 **Median filter:** A median filter replaces each pixel by the median value of its neighbors. This is especially useful for removing salt- and-pepper noise, where the noise takes the form of some pixels having very high or very low values of intensity.

3 **Convolution:** The convolution operation takes an image and a filter and produces a new image. The filter is represented as a 2D matrix of size k x k. The convolution operation then takes k x k neighborhoods in the image, multiplies it element-wise with the filter, and sums up the result to produce the output value for the center pixel.

4 **Box filter:** A k x k box filter is a filter filled with a constant value of 1/k^2^. Convolving with this filter replaces every pixel by the average of its neighbors in a k x k neighborhood.

5 **Connected component analysis:** Given a binary image, connected component analysis identifies contiguous regions of the image which have a ‘True’ value. This analysis is done by interpreting the image as a graph where True pixels are nodes and there are edges between every pair of neighboring pixels. Connected components in this graph correspond to contiguous regions in the image.

